# Tracheal tube fusion in *Drosophila* involves release of extracellular vesicles from multivesicular bodies

**DOI:** 10.1101/2021.08.22.457102

**Authors:** Carolina Camelo, Anna Körte, Thea Jacobs, Stefan Luschnig

## Abstract

Extracellular vesicles (EVs) comprise diverse types of cell-released membranous structures that are thought to play important roles in intercellular communication. Despite extensive work on the formation and functions of EVs in cultured cells, studies of EVs *in vivo* have remained scarce. We report here that EVs are present in the developing lumen of tracheal tubes in *Drosophila* embryos. We defined two distinct EV subpopulations, one of which contains the Munc13-4 homologue Staccato (Stac) and is spatially and temporally associated with tracheal tube fusion events. The formation of Stac-positive luminal EVs depends on the tip-cell-specific GTPase Arl3, which is also required for the formation of Stac-positive multivesicular bodies, suggesting that Stac-EVs derive from fusion of Stac-MVBs with the luminal membrane in tip cells during anastomosis formation. The GTPases Rab27 and Rab35 cooperate downstream of Arl3 to promote Stac-MVB formation and tube fusion. We propose that Stac-MVBs act as membrane reservoirs that facilitate tracheal lumen fusion in a process regulated by Arl3, Rab27, Rab35, and Stac/Munc13-4.

## Introduction

Tubular organs, such as the vertebrate vasculature or the *Drosophila* tracheae, develop from separate units that fuse to form tubular networks. Tracheal tube fusion in *Drosophila* is mediated by specialized ‘fusion’ cells (FCs; Samakovlis *et al*., 1996) that transform into lumenized toroids to connect adjacent tubes. The fusion of luminal membranes inside FCs depends on the Munc13-4 orthologue Staccato (Stac; Caviglia et al., 2016), which localizes to lysosome-related organelles (LROs) specifically in FCs. Munc13-4 acts as an essential priming factor for SNARE complex formation and SNARE-mediated membrane fusion (Boswell et al., 2012) and interacts with the GTPase Rab27 to promote membrane docking and exocytosis of LROs in hematopoietic cells, melanocytes, and cancer cells (Alzahofi *et al*., 2020; Johnson *et al*., 2011; Messenger *et al*., 2018; Neeft *et al*., 2005; Shirakawa *et al*., 2004). Formation of Stac-positive LROs in FCs depends on the GTPase Arl3 (Caviglia *et al*., 2016), which is specifically expressed in FCs and was proposed to regulate targeting of the exocytosis machinery to the apical plasma membrane (Jiang *et al*., 2007; Kakihara *et al*., 2008). However, how LROs mediate tracheal tube fusion was not clear.

Recent work showed that Munc13-4 regulates multivesicular body (MVB) maturation and release of exosomes in cultured cancer cells (Messenger et al., 2018). Exosomes originate from exocytosis of endolysosomal MVBs, while ectosomes, another class of membranous extracellular vesicles (EVs), bud directly from the plasma membrane (Kalluri and LeBleu, 2020). EVs are thought to play important roles in intercellular communication during development, homeostasis and pathological conditions. However, despite many studies on the biogenesis and functions of EVs in cultured cells, evidence of EVs and their physiological functions *in vivo* has remained rare (Corrigan et al., 2014; Scott et al., 2021; Tsai et al., 2019; Verweij et al., 2018).

We report here the discovery of EVs in the developing tracheal lumen of *Drosophila* embryos. We defined two distinct EV subpopulations, one of which contains Stac/Munc13-4 and is spatially and temporally associated with tracheal tube fusion events. The presence of Stac-EVs depends on *Arl3* function in tracheal FCs. We identified Rab GTPases required for Stac-MVB formation and show that Rab27 and Rab35 act in a partially redundant fashion downstream of Arl3 to promote Stac-MVB formation and tracheal tube fusion. We propose that Stac-positive EVs originate from the fusion of multivesicular LROs with the luminal plasma membrane during anastomosis formation in FCs.

## Results

### Distinct populations of extracellular vesicles are present in the embryonic tracheal lumen

When expressed in the embryonic tracheal system, EGFP-Stac and mCherry-Stac were distributed throughout the cytoplasm of tracheal cells, while only in FCs the proteins accumulated at LROs (Fig. 1A; Caviglia *et al*., 2016). Surprisingly, we noticed EGFP-Stac- or mCherry-Stac-positive puncta also inside the tracheal lumen of embryos at stage 15 (Fig. 1A-C; Movie S1). Extracellular EGFP-Stac puncta were submicron-sized (0.60 µm +/−0.12 µm diameter based on confocal microscopy; n=19 EVs; Fig. 1H) and were surrounded by luminal material containing the secreted protein Vermiform-mRFP (Verm-mRFP; Fig. 1A). EGFP-Stac puncta were detectable in the lumen of the dorsal trunk (DT) tube in 40% of the embryos (Fig. 1G; n=50 embryos). The puncta were labeled by the membrane marker palmitoylated mKate2 (palm-mKate2; Fig. 1B), suggesting that the luminal Stac puncta are membranous extracellular vesicles (EVs). Stac puncta were also labeled by CD63-GFP, a GFP-tagged vertebrate tetraspanin that localizes to the limiting membrane and to membranes of internal vesicles of multivesicular bodies (MVBs) and is released on exosomes in mammalian tissue culture as well as in *Drosophila* (Fig. 1C; Escola *et al*., 1998; Panáková *et al*., 2005). Interestingly, we observed a larger pool of CD63-GFP-positive EVs, only a fraction (2%, n=89 EVs in 11 embryos) of which were also positive for mCherry-Stac (Fig. 1C). EVs marked by CD63-GFP, but not by mCherry-Stac, were found in all embryos analyzed (n=14; Fig. 1C,G) and showed an apparent size similar to EGFP-Stac-positive EVs (diameter 0,63 µm +/−0,25 µm; n=79 EVs in 11 embryos; Fig. 1H). These findings indicate that the embryonic tracheal lumen contains distinct pools of EVs, a subset of which accumulates Munc13-4/Stac.

**Figure 1.**
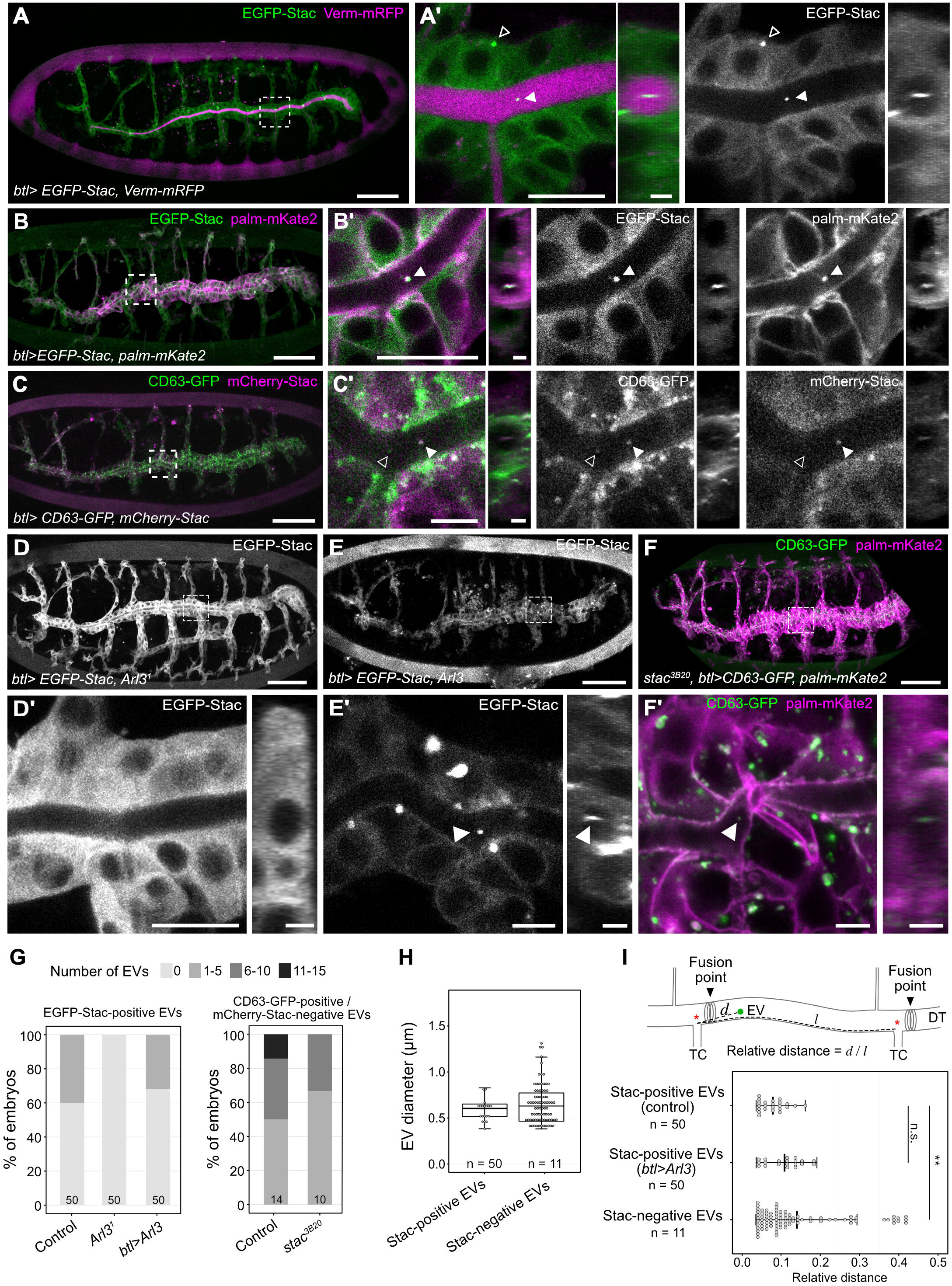
Extracellular vesicles are present in the embryonic tracheal lumen. **(A-B)** Lateral view of living embryos (stage 15) expressing EGFP-Stac (green) and Vermiform-mRFP (Verm-mRFP, magenta; A) or EGFP-Stac (green) and palmitoylated mKate2 (palm-mKate2, magenta; B) in tracheal cells under the control of *btl-Gal4*. (A’, B’) are close-ups of the regions indicated in (A, B). EGFP-Stac is distributed throughout the cytosol and accumulates at LROs (open arrowheads) only in tracheal fusion cells. Note Stac-positive EVs (filled arrowheads) in the dorsal trunk lumen. Owing to limited axial resolution, EVs could not be identified unambiguously in the smaller-caliber lumina of other tracheal branches. **(C)** Lateral view of living embryos (stage 15) expressing CD63-GFP (green) and mCherry-Stac (magenta) in tracheal cells. CD63-GFP labels mCherry-Stac-positive EVs (filled arrowhead) and mCherry-Stac-negative EVs (open arrowhead). **(D-E)** Lateral view of living *Arl3*^*1*^ embryo (stage 15) expressing EGFP-Stac in tracheal cells in (D) and embryo co-expressing EGFP-Stac and Arl3 in tracheal cells (E). (D’ and E’) are close-ups indicated in (D, E). Note that Stac-LROs and Stac-EVs are absent in *Arl3*^*1*^ mutant (D), whereas Stac-LROs are induced ectopically upon Arl3 misexpression (E). **(F)** Lateral view of living *stac*^*3B20*^ embryo (stage 15) expressing CD63-GFP (green) and palm-mKate2 (magenta) in tracheal cells. (F’) shows close-up of region indicated in (F). **(G)** Quantification of EGFP-Stac-positive (left) or CD63-GFP-positive / mCherry-Stac-negative (right) EVs. Number of tracheal EVs per embryo was determined in the indicated genotypes. Embryos expressing EGFP-Stac (left) or CD63-GFP and mCherry-Stac <u>(right)</u> under the control of *btl*-Gal4 in tracheal cells were used as controls. Number of embryos analyzed (n) is indicated for each genotype. **(H)** Diameter of Stac-positive and Stac-negative EVs as detected by confocal microscopy. Each data point in the graph represents one EV. Number of embryos (n) is indicated. **(I)** Quantification of the relative distance of EVs to the closest tracheal fusion point (marked by adjacent transverse connective (TC) branch; asterisk) in the DT tube. Each data point represents one EV. Mean +/− SD and the number of embryos scored (n) are indicated. T-test: ** p < 0.001; ns, not significant. Scale bars: (A-F) 50 µm; (A’, B’, C’, D’, E’, F’) 10 µm; cross-sections 2 µm.

### Stac-positive EVs are associated with tracheal tube fusion sites

Luminal EVs were largely immobile over the course of 240 minutes (Fig. S1A,B, Movie S2, Movie S3), suggesting that EVs are trapped in the fibrillar matrix that fills the embryonic tracheal lumen (Dong *et al*., 2014; Tonning *et al*., 2006). The EVs subsequently disappeared during tracheal liquid clearance (Fig. S1C, Movie S3). Interestingly, Stac-positive EVs showed a non-uniform distribution along the tracheal lumen and were frequently found close to tube fusion sites at tracheal metamere boundaries (marked by the transverse connective branch; Fig. 1I, Fig. S1C), suggesting that Stac-positive EVs might be released by FCs during tube fusion. While direct visualization of tracheal EV exocytosis events was precluded by the small diameter of the tracheal lumen during tube fusion and by the small size of EVs, we were able to trace luminal EVs in time-lapse movies to DT fusion points (Fig. S1, Movie S2), suggesting that Stac-positive EVs are released during the tube fusion process.

### Formation of Stac-positive EVs, but not of EVs in general, depends on the GTPase Arl3

Consistent with the notion that Stac-positive EVs are released during tube fusion, we did not detect Stac-positive EVs in the tracheal lumen of *Arl3*^*1*^ embryos (n=50), which lack Stac-LROs in FCs (Fig. 1D,G; Caviglia et al., 2016), indicating that the FC-specific GTPase Arl3 is required for Stac-EV formation, and corroborating the notion that Stac-EVs originate from Stac-LROs in FCs. Conversely, to ask whether the presence of Stac-LROs is sufficient to enable tracheal cells to release Stac-EVs, we generated ectopic Stac-LROs throughout the tracheal system by mis-expressing Arl3 in all tracheal cells (Fig. 1E; Caviglia *et al*., 2016). However, Arl3 misexpression did not affect the number (Fig. 1E,G) or spatial distribution (Fig. 1I) of Stac-EVs in the DT lumen, suggesting that Stac-EVs are released exclusively by FCs at tracheal anastomosis sites, even when Stac-LROs are ectopically induced in other tracheal cells.

By contrast, CD63-GFP-positive / mCherry-Stac-negative EVs were distributed more broadly throughout the lumen (Fig. 1I). CD63-GFP-positive EVs were present in the tracheal lumen of tube-fusion-defective *stac*^*3B20*^ embryos, albeit at reduced numbers compared to controls (Fig. 1F,G; n=10 embryos), indicating that the formation of CD63-GFP-positive EVs does not generally depend on tube fusion or *stac* function. Taken together, these findings suggest that CD63-GFP-positive/Stac-negative EVs are produced by all tracheal cells, while Stac-positive EVs are associated with FCs and their formation depends on the FC-specific GTPase Arl3.

### Stac-LROs display features of multivesicular bodies

Based on these results, we hypothesized that Stac-positive EVs might originate from multivesicular membrane compartments in FCs. Supporting this idea, Stac-LROs were labeled by the MVB marker CD63-GFP (Fig. 2A) and mCherry-Stac was accumulating inside and at the boundary of CD63-GFP-limited vesicular structures (Fig. 2B), suggesting that mCherry-Stac is present in intraluminal vesicles (ILVs) and at the outer membrane of MVBs (Fig. 2B’). To test whether Stac protein can be incorporated into ILVs in a well-characterized system for MVB and exosome biology, we used *Abd-B*-Gal4 to express EGFP-Stac in secondary cells of the adult testis accessory gland. These cells contain large endolysosomal multivesicular compartments and release EVs into the accessory gland lumen (Fig. S2; Corrigan *et al*., 2014; Fan *et al*., 2020). Indeed, EGFP-Stac was present in ILVs inside MVBs in secondary cells, as well as in small membranous EVs in the accessory gland lumen (Fig. S2D-E). Thus, Stac can be incorporated into ILVs and can be secreted as ILV-derived EVs into the accessory gland and the tracheal lumen. Consistent with the hypothesis that Stac-MVBs secrete EVs during tube fusion, we observed Stac-MVBs between the tips of approaching FC lumina (labeled by the apical membrane marker PLC δ-PH-EGFP; Fig. S3D), and these MVBs appeared to fuse with the central lumen at the FC-FC interface (labeled by the adherens junction component E-Cad::3xmTagRFP; Fig. S3B).

**Figure 2.**
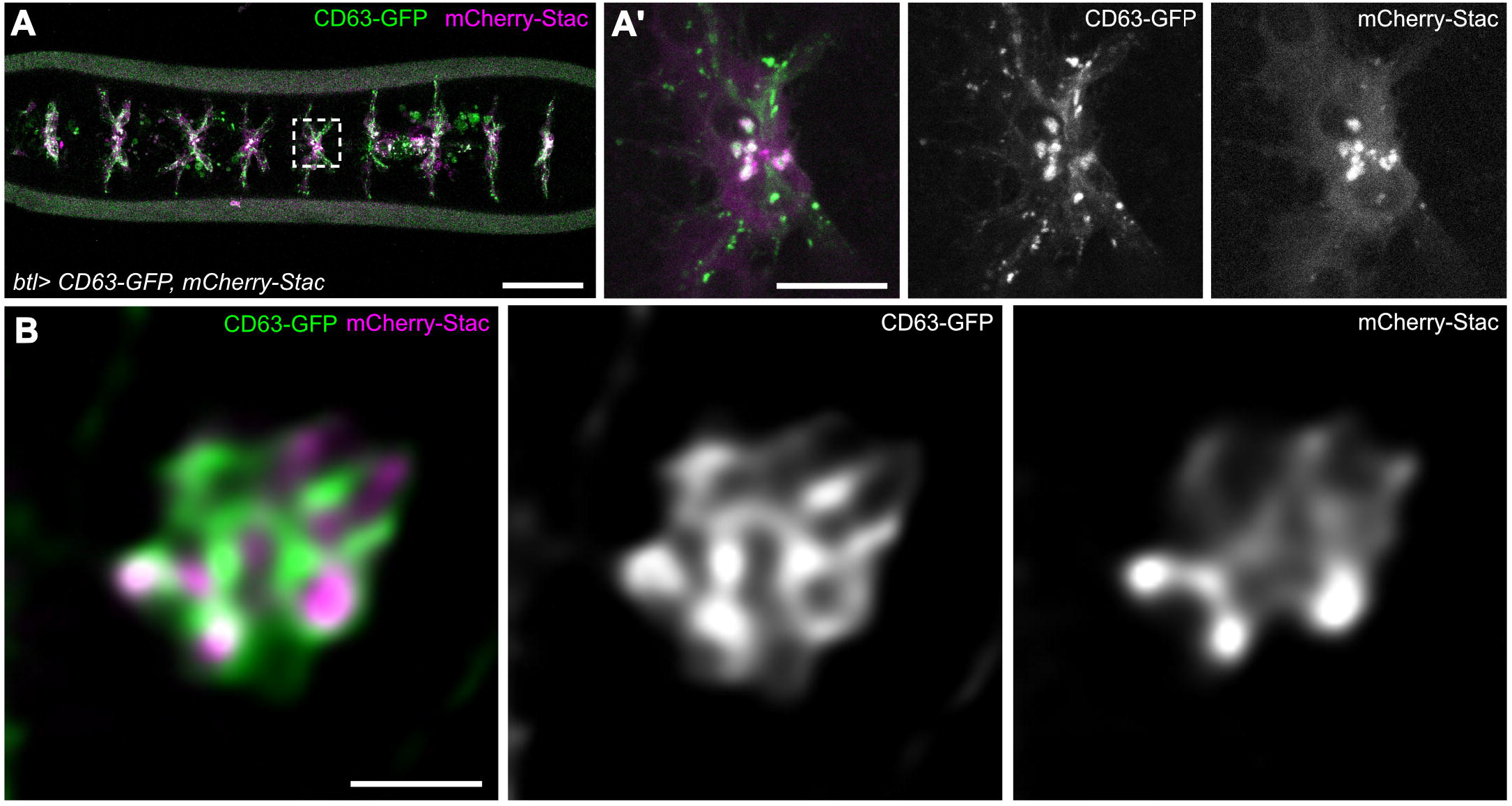
Stac-MVBs colocalize with CD63-GFP and localize between invading lumina. **(A)** Dorsal view of living embryo (stage 15) expressing the MVB marker CD63-GFP (green) and mCherry-Stac (magenta) in tracheal cells. Note that mCherry-Stac colocalizes with CD63-GFP (arrowheads). (A’) shows close-up of region indicated in (A). **(B)** Close-up of mCherry-Stac puncta. Note that mCherry-Stac signals are bounded by CD63-GFP. The image was processed using deconvolution. Scale bars: (A) 50 µm; (A’) 10 µm; (B) 1 µm.

### Stac and Arl3 function together and co-localize in tracheal cells

The GTPase Arl3 is necessary and sufficient for Stac-LRO formation in tracheal cells (Caviglia *et al*., 2016), and is required for the presence of Stac-EVs in the tracheal lumen (Fig. 1D,G). Arl3 may act by modulating the activity of Rab GTPases that recruit Stac to LROs, in analogy to the proposed role of Arl3 in mammalian cells (Ismail, 2011; Williams, 2011). To test whether Arl3 and Stac act in parallel or in the same pathway to promote Stac-LRO formation and tube fusion, we analyzed tracheal defects in amorphic *Arl3*^*1*^ and *stac*^*3B20*^ single mutant and in *Arl3*^*1*^ *stac*^*3B20*^ double mutant embryos (Fig. 3A-E). *Arl3*^*1*^ and *stac*^*3B20*^ embryos displayed tube fusion defects of similar expressivity, with on average two out of nine fused DT metamere anastomoses (Fig. 3E). Fusion defects were not enhanced in *Arl3*^*1*^ *stac*^*3B20*^ double mutants (Fig. 3D,E), suggesting that Arl3 and Stac do not function independently, but act in the same pathway. Consistent with this notion, an HA-epitope-tagged Arl3 protein (Arl3-HA) colocalized with a subset of EGFP-Stac puncta when the two proteins were expressed in tracheal cells (Fig. 3F). These results suggest that Arl3 and Stac function in a common pathway to mediate the formation of Stac-MVBs.

**Figure 3.**
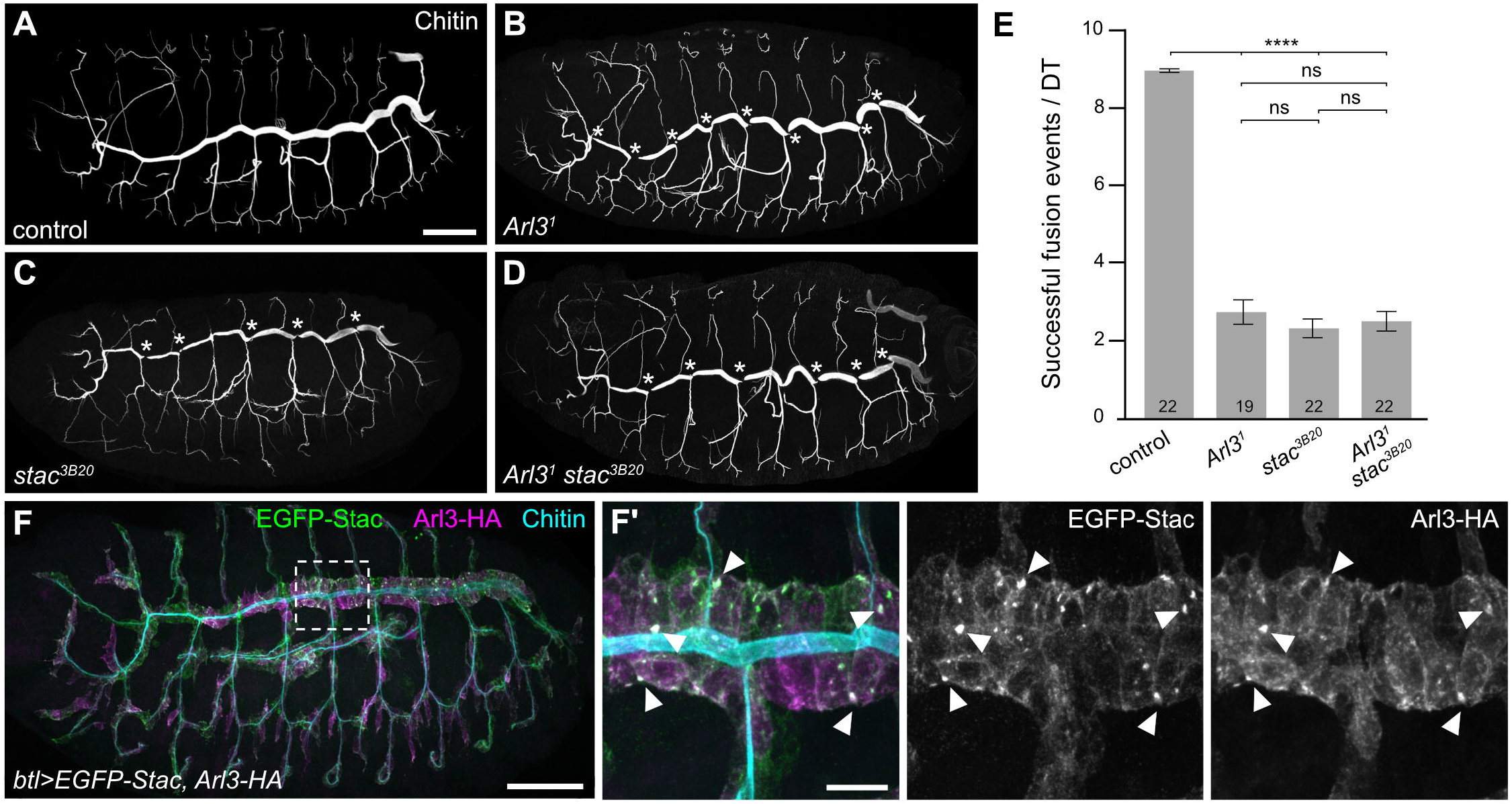
Arl3 and Stac act in the same pathway and co-localize in tracheal cells. **(A-D)** Lateral views of control (*y w*; A), *Arl3*^*1*^ (B), *stac*^*3B20*^ (C) and *Arl3*^*1*^ *stac*^*3B20*^ (D) double mutant embryos (stage 15) stained for chitin to label tracheal lumen. Asterisks indicate DT tube fusion defects. **(E)** Quantification of DT fusion defects. Control (*y w*) embryos show nine anastomoses (each indicating a successful fusion event) per DT tube. Note that *Arl3*^*1*^ and *stac*^*3B20*^ single mutants show DT fusion defects of similar extent as in *Arl3*^*1*^ *stac*^*3B20*^ double mutants. Number of embryos (n) is indicated. Wilcoxon rank-sum test: **** p < 0.0001; ns, not significant. **(F)** Lateral view of embryo expressing EGFP-Stac (green) and Arl3-HA (magenta) in tracheal cells. Chitin (cyan) labels tracheal lumen. Note that Arl3-HA colocalizes with a subset of EGFP-Stac puncta (arrowheads). Scale bars: (A,B,C,D,F) 50 µm; (F’) 10 µm.

### Identification of Rab GTPases involved in Stac-LRO formation

To identify factors acting downstream of Arl3 to promote Stac-LRO formation, we generated a sensitized genetic background by mis-expressing Arl3 throughout the tracheal system in EGFP-Stac-expressing embryos. This led to ectopic Stac-LROs in most tracheal cells (44+/−24 Stac-LROs in DT metameres 4-6, n=10; Fig. S4), whereas Stac-LROs were restricted to FCs in wild-type embryos (Fig. 1A). To identify Rab GTPases required for Stac-LRO formation, we reduced in this genetic background the gene dosage of each of 26 of the 33 annotated *Drosophila* Rab GTPases by introducing null mutations of the respective *Rab* loci (Chan *et al*., 2011; Kohrs *et al*., 2021). We analyzed the resulting *Rab* heterozygous or hemizygous embryos by determining the number of Stac-LROs (Fig. S4). This approach revealed six Rab GTPases (*Rab8, Rab10, Rab27, Rab30, Rab35, Rab39*) whose reduced gene dosage led to significantly reduced Stac-LRO numbers compared to control embryos (Fig. S4B). We focused on Rab27, Rab35, and Rab39, which are strong candidates for regulators of late-endosomal trafficking, LRO/MVB membrane docking (Alzahofi *et al*., 2020; Biesemann *et al*., 2017; Caviglia *et al*., 2016; Neeft *et al*., 2005), or exosome release (Messenger *et al*., 2018), whereas Rab8, Rab10 and Rab30 are predicted to play more general roles in ER morphology, Golgi sorting, and retrograde transport, respectively (Bellec *et al*., 2018; English and Voeltz, 2013; Gillingham *et al*., 2014; Schuck *et al*., 2007).

### Rab27 and Rab35 cooperate to promote Stac-LRO formation

Having identified *Rab27, Rab35* and *Rab39* mutations as suppressors of ectopic Stac-LRO formation upon Arl3 misexpression, we tested whether these Rab GTPases are also required for the formation of entopic Stac-LROs in FCs of otherwise wild-type embryos. Embryos lacking maternal and zygotic Rab27 or zygotic Rab35 displayed significantly reduced numbers of Stac-LRO in FCs (47% and 53%, respectively, compared to control; Fig. 4A,C,D,F), suggesting that these GTPases participate in Stac-LRO formation. Surprisingly, however, the number of Stac-LROs in *Rab39^−^* embryos was not different from control embryos (Fig. 4A,B,F), despite the suppression of ectopic Stac-LROs in Arl3-misexpressing embryos (Fig. S4), suggesting that Rab39 is not essential for forming Stac-LROs in FCs. Because loss of Rab27 or of Rab35 each caused a partial reduction of Stac-LROs, we generated a *Rab27^−^ Rab35^−^* double mutant to ask whether Rab27 and Rab35 cooperate to form Stac-LROs. Indeed, *Rab27^−^ Rab35^−^* embryos showed significantly fewer Stac-LROs compared to *Rab27^−^* and *Rab35^−^* single mutants (30.5% and 27%, respectively; Fig. 4A,C-F), suggesting that Rab27 and Rab35 cooperate in Stac-LRO formation. Consistent with these findings, endogenously tagged YFP::Rab35 (Fig. 4M; Caviglia *et al*., 2016) and overexpressed YFP-Rab27 (Fig. 4N) overlapped with a subset of mCherry-Stac-LROs in FCs. Moreover, *Rab27^−^* and *Rab35^−^* embryos exhibited slightly reduced numbers of Stac-EVs in the tracheal lumen. 80% of *Rab27^−^* and 76% of *Rab35^−^* embryos showed no EVs, whereas at least one EV was detectable in 40% of control embryos (n=50 embryos per genotype). Taken together, these results suggest that Rab27 and Rab35 cooperate to promote the formation of Stac-LROs and possibly the release of Stac-EVs.

**Figure 4.**
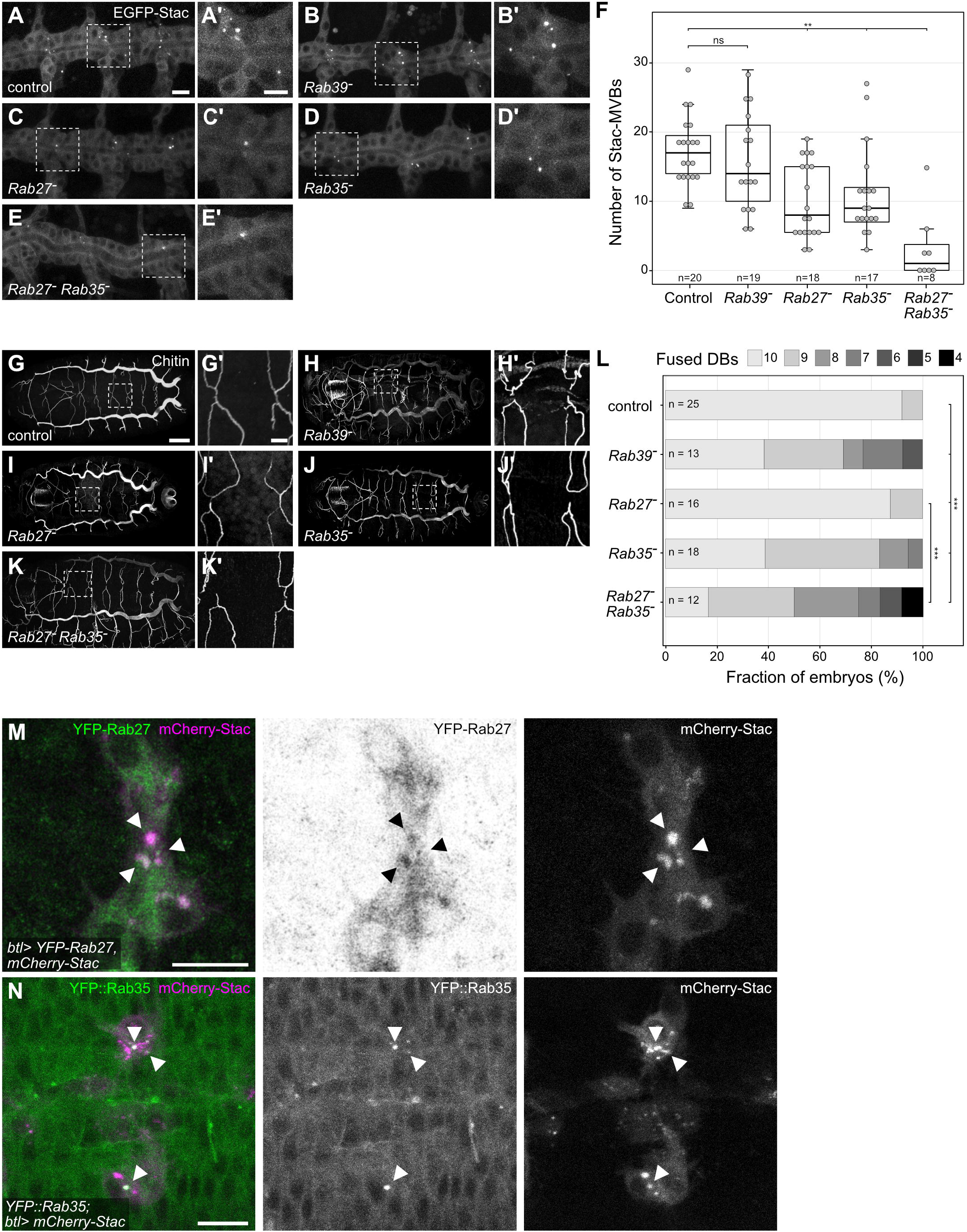
Rab27 and Rab35 act downstream of Arl3 to regulate Stac-MVB formation. **(A-E)** Lateral views of embryos (stage 15) of the indicated genotypes expressing EGFP-Stac in tracheal cells under the control of *btl-Gal4*. (A’-D’) are close-ups indicated in (A-D). Note the presence of EGFP-Stac-MVBs in tracheal fusion cells. *btl-Gal4 UAS-EGFP-Stac* embryos were used as control. **(F)** Quantification of EGFP-Stac-MVBs in DT metameres 4-6. Genotypes and number of embryos scored (n) are indicated. *btl-Gal4 UAS-EGFP-Stac* embryos were used as control. Wilcoxon rank-sum test: ** p < 0.001; ns, not significant. **(G-K)** Dorsal views of embryos (stage 16) of the indicated genotypes. Chitin staining labels the tracheal lumen. (F’-J’) are close-ups indicated in (F-J). *y w* embryos were used as control. **(L)** Quantification of DB fusion defects. Number of fused DBs per embryo is indicated by different shades of grey. Genotypes and numbers of embryos (n) analyzed are indicated. *y w* embryos were used as control. Permutation statistical test: *** p < 0.0001. **(M)** DB FCs of embryo expressing mCherry-Stac (magenta) and YFP-Rab27 (green) under the control of *btl-Gal4*. **(N)** DB FCs of embryo expressing mCherry-Stac (magenta) under the control of *btl-Gal4* and endogenously tagged YFP::Rab35 (green). Scale bars: (A,B,C,D,E, A’,B’,C’,D’,E’) 10 µm; (G,H,I,J,K) 50 µm; (G’,H’,I’,J’,K’), 10 µm; (M,N) 10 µm.

### Rab27, Rab35 and Rab39 are required for tracheal tube fusion

We finally asked whether Rab27, Rab35, and Rab39 are required for tracheal tube fusion. While *Rab27^−^* and *Rab39^−^* homozygous (female) and hemizygous (male) flies were viable and fertile, *Rab35^−^* flies were semi-lethal, with most animals dying during the pupal stage, and surviving adult *Rab35^−^* males were sterile. Embryos lacking maternal and zygotic Rab27 or Rab39 and embryos lacking zygotic Rab35 developed normally and did not show DT fusion defects (Fig. 4A-E, Fig. 4G-K). To assess more subtle tracheal defects in the mutant embryos, we analyzed dorsal branch (DB) fusion, which is more sensitive towards genetic perturbations than DT fusion. While nearly all ten DBs per embryo were fused in controls (Fig. 4G,L; average of 9.9 fused DBs, n=25 embryos), *Rab35^−^* (n=18) and *Rab39^−^* (n=13) embryos showed on average 9.1 and 8.8 fused DBs, respectively, per embryo (Fig. 4G,H,J,L). By contrast, *Rab27^−^* embryos did not show notable tracheal defects (n=16; Fig. 4I,L). However, double mutant embryos lacking zygotic Rab27 and Rab35 showed enhanced DB fusion defects (average of 8.1 fused DBs per embryo; n=12) compared to *Rab27^−^* and *Rab35^−^* single mutants (Fig. 4K,L), indicating that Rab27 and Rab35 carry out partially redundant functions in tracheal tube fusion.

## Discussion

We report the presence of membranous EVs in the lumen of the developing tracheal system in *Drosophila* embryos. Intriguingly, a subset of these luminal EVs is associated with tracheal tube fusion sites and carries the Munc13-4 orthologue Stac. In tracheal fusion cells, Stac associates with LROs that display features of multivesicular membrane compartments. We show that the formation of these compartments and of luminal-Stac EVs depends on the FC-specific GTPase Arl3, suggesting that Stac-EVs originate from fusion of Stac-MVBs with the luminal plasma membrane in tracheal tip cells. A systematic survey of Rab GTPases revealed a set of Rab proteins required for Stac-MVB formation in tracheal cells. We showed that Rab27 and Rab35 act in a partially redundant manner downstream of Arl3 to promote Stac-MVB formation and tracheal tube fusion. These findings suggest that tracheal anastomosis formation proceeds through fusion of Stac-MVBs with the luminal plasma membrane in tracheal tip cells, resulting in release of Stac-positive EVs. Our attempts to directly visualize tracheal EV exocytosis events using the pH-sensitive fluorescent proteins pHluorin (Sankaranarayanan *et al*., 2000) or pHuji (Shen *et al*., 2014) fused to the lysosomal transmembrane protein Lamp1 were not successful (not shown). However, we were able to detect EVs that emerged at and remained associated with anastomosis sites upon completion of DT fusion, supporting the idea that Stac-positive EVs are released during tube fusion. Moreover, we showed that Stac protein is incorporated into ILVs and is secreted in exosomes when expressed in male accessory gland secondary cells, a model system for exosome biology in *Drosophila*.

How is the formation of MVBs in FCs regulated? We found that Rab27 and Rab35 act downstream of Arl3 in a cooperative fashion to promote Stac-MVB formation. Consistent with our findings in *Drosophila*, Rab27a and Rab27b (Ostrowski *et al*., 2010) and Rab35 (Hsu *et al*., 2010) regulate exosome secretion in mammalian cells. Moreover, the Stac homologue Munc13-4 is a key effector of Rab27a in LRO docking in hematopoietic cells and melanocytes (Alzahofi *et al*., 2020; Johnson *et al*., 2011; Neeft *et al*., 2005; Shirakawa *et al*., 2004), and regulates MVB maturation and exosome release in cancer cells (Messenger *et al*., 2018). Whereas Rab27 and Rab35 are required for Stac-MVB formation or for anchoring Stac to these compartments, Rab39 is dispensable for FC-specific Stac-LRO formation, suggesting that it acts in a successive step, such as intracellular transport or plasma membrane docking of LROs. While our findings define a core pathway of MVB maturation and EV release in tracheal tip cells, additional players, *e*.*g*., components of the ESCRT complex, are likely to participate in MVB formation. MVBs were identified and proposed to act as intermediates in intracellular lumen formation also in tracheal terminal cells, which like fusion cells form seamless tubes (Mathew *et al*., 2020; Nikolova and Metzstein, 2015). Interestingly, Rab35 directs polarized transport of apical components to promote terminal cell lumen growth (Schottenfeld-Roames and Ghabrial, 2012). Whether Rab35 participates in MVB formation also in tracheal terminal cells remains to be investigated.

Extracellular vesicles were studied extensively in the context of intercellular communication and were proposed to play important roles in organ homeostasis and pathology. However, despite substantial progress in characterizing EVs and their mode of action in cultured cells, studies of physiological functions of EVs *in vivo* have remained scarce. For instance, exosomes are released into the seminal fluid in *Drosophila* male accessory glands and modulate female reproductive behavior (Corrigan *et al*., 2014). Glia-derived exosomes stimulate growth of motor neurons and tracheae in *Drosophila* larvae through exosomal microRNA-dependent gene regulation in target cells (Tsai *et al*., 2019). Moreover, work in zebrafish revealed that cardiomyocyte-derived EVs are taken up by macrophages and endothelial cells (Scott *et al*., 2021), and exosomes were proposed to play a role in trophic support of endothelial cells during vasculogenesis (Verweij *et al*., 2018). It will be exciting to explore possible functions of tracheal EVs, *e*.*g*. in mediating long-range cell-to-cell communication across the tracheal lumen. However, it is unlikely that the luminal EVs that we discovered in late-stage embryos participate in such intercellular communication, as the presence at this stage of a luminal matrix (Dong *et al*., 2014; Tonning *et al*., 2006) appears to constrain EV mobility, and the cuticle covering the apical surface of tracheal cells presumably prevents fusion of luminal EVs with the plasma membrane. Rather, Stac-positive EVs are likely to constitute remnants of MVB-plasma membrane fusion events during anastomosis formation. Stac-MVBs might serve as membrane reservoirs that facilitate the final step of lumen fusion. ILVs might back-fuse with the MVB limiting membrane, resulting in a rapid increase of membrane material available at the luminal membrane interfaces in FCs (Figure S5). Such a mechanism was described in dendritic cells, where MVBs carrying major histocompatibility complex class II (MHC II) in ILVs are reorganized after stimulation, resulting in long tubular membrane extensions and transport of MHCII to the plasma membrane (Kleijmeer *et al*., 2001). Recruitment of ILVs to the MVB limiting membrane will increase the surface area, but may also lead to surface exposure of ILV membrane proteins that could promote membrane fusion. How MVB exocytosis and EV release are regulated in tracheal cells, and whether related mechanisms are involved in anastomosis formation in vertebrates, will be exciting subjects for future studies.

## Acknowledgements

We thank Wilko Backer for expert technical help, Raphael Schleutker for help with image analysis and statistics, and Yohanns Bellaiche, Marko Brankatschk, Natalie Dye, Suzanne Eaton, Shigeo Hayashi, Robin Hiesinger, and Clive Wilson for providing fly stocks and reagents. We thank Sara Caviglia, Mylène Lancino and Raphael Schleutker for comments on the manuscript. Work in the Luschnig laboratory was supported by the Deutsche Forschungsgemeinschaft (SFB 1348 “Dynamic Cellular Interfaces”; SFB 1009 “Breaking Barriers”), the “Cells-in-Motion” Cluster of Excellence (EXC 1003-CiM) and the University of Münster.

## Author contributions

Conceptualization, all authors; Methodology, all authors; Investigation, C.C., A.K., T.J.; Formal Analysis, C.C., A.K., T.J.; Visualization, C.C., A.K., T.J.; Reagents and tools, all authors; Writing – Original Draft, C.C.; Writing – Review and Editing, C.C., S.L.; Funding Acquisition, S.L; Supervision, S.L.

## Materials and Methods

### *Drosophila* strains and genetics

The following *Drosophila* strains are described in FlyBase: *Abd-B-Gal4, btl-Gal4, UAS-PLC* δ*-PH-EGFP, UAS-EGFP-Stac, UAS-mCherry-Stac, UAS-Verm-mRFP, UAS-palm-mKate2* (Caviglia *et al*., 2016), *UAS-CD63-GFP* (Panáková *et al*., 2005; gift from S. Eaton), *E-Cad::3xmTagRFP* (Pinheiro *et al*., 2017; gift from Y. Bellaiche), *Arl3*^*1*^, *UAS-Arl3* (Kakihara *et al*., 2008; gift from S. Hayashi), *UAS-Arl3-3xHA* (Bischof *et al*., 2007), *stac*^*3B20*^ (Caviglia *et al*., 2016), *UAS-YFP-Rab27* (Zhang *et al*., 2007), *YFP::Rab35* (Dunst *et al*., 2015). The collection on *Rab* null alleles are described in (Chan *et al*., 2011; Kohrs *et al*., 2021). Lethal *Rab* mutants were balanced using *FM7 Dfd-GMR-nvYFP, CyO Dfd-GMR-nvYFP*, or *TM6B Dfd-GMR-nvYFP* balancer chromosomes (Le *et al*., 2006). For analyzing the effects of autosomal *Rab* mutations on Stac-LRO number, *btl-Gal4 UAS-EGFP-Stac* females were crossed to males carrying the *Rab^−^* mutation. For X-chromosomal *Rab* loci, females carrying the *Rab^−^* mutation were crossed to *btl-Gal4 UAS-EGFP-Stac* males. Embryos (14-18 h) were collected at 22°C. Embryos carrying the *Rab^−^* allele were identified by the absence of *Dfd*-YFP expression. The *y*^*1*^ *w*^*1118*^ strain was used as wild-type control.

### Constructs and transgenic flies

UAS-OLLAS-pHluorin-Lamp1 and UAS-OLLAS-pHuji-Lamp1 constructs were generated by fusing the OLLAS-pHluorin (Sankaranarayanan *et al*., 2000) or OLLAS-pHuji (Shen *et al*., 2014) ORFs to an N-terminal preprolactin signal peptide and C-terminally to the transmembrane domain and cytosolic domain of the human lysosomal transmembrane protein Lamp1 (Pulipparacharuvil *et al*., 2005), which localizes to Stac-LROs in FCs (Caviglia *et al*., 2016). The constructs were inserted into the pUASt-attB vector (using EcoRI and XbaI restriction sites) and and integrated into the *attP40* and *attP2* landing sites using PhiC31 integrase (Bischof *et al*., 2007).

### Antibodies and immunostaining

Embryos were collected 14-18 hours after egg lay (h AEL) and 17h-18 h AEL (22°C) for analysis of DT and DB fusion defects, respectively. Embryos were fixed in 4 % formaldehyde in PBS/heptane for 20 min and devitellinized by shaking in methanol/heptane. The following primary antibodies were used: mouse anti-GFP (1:500; Sigma G6539), chicken anti-GFP (1:500; Abcam 13970), rat anti-HA (1:300; Roche 3F10). Luminal chitin was detected using the chitin-binding domain from *Bacillus circulans* chitinase A1 conjugated with SNAP-Surface AlexaFluor 488 or 563, produced as described previously (Caviglia and Luschnig, 2013).

### Microscopy and image analysis

For live imaging, dechorionated embryos were glued on a coverslip (0.17 mm, grade #1.5), embedded in Voltalef 10S oil and covered with a gas-permeable foil (Lumox, Sarstedt). Accessory glands were dissected from three-day-old virgin males, stained with membrane dye MM4-64 (2 nM; Santa Cruz Biotechnology) and Hoechst 33342 (1 μg/ml; Sigma) to label plasma membrane and nuclei, respectively, and were mounted on slides in cold (4°C) PBS. Imaging was performed on a Leica SP8 confocal microscope with 40x/1.3 NA and 63x/1.4 NA objectives and HyD detectors or on a Zeiss LSM710 confocal microscope with a 40x/1.1 NA objective. Images were processed using OMERO (5.4.10), Fiji (ImageJ; 1.53c), and Imaris (8.4.1, Bitplane), and were prepared in Affinity Designer (1.8.4). Cross-sections were performed using Imaris (8.4.1). Where indicated, images were deconvolved using the Leica HyVolution software in adaptive mode. Calculations were performed in R (3.5.1) using RStudio Interface (1.3.1093).

### Quantification of EV number, diameter and distance from tracheal fusion points

Confocal Z-stacks of living embryos (stages 15 and 16) were analyzed for intraluminal signals of EGFP-Stac, mCherry-Stac, or CD63-GFP. For measuring EV diameter, a single slice containing the EV was analyzed. EVs were segmented by applying a median filter (radius = 2) and using the Automatic Fiji threshold Intermodes. Feret’s diameter of the vesicles was measured using the Analyze Particles plugin in Fiji. For measuring the distance of EVs from the closest DT anastomosis, the base of the closest transverse connective (TC) branch was used as a morphological landmark adjacent to the DT fusion point. The distance between the EV and the closest TC (*d*) and the length (*l*) of the corresponding DT metamere (distance between the two flanking TCs) were measured in 3D using Imaris (8.4.1). The relative distance was calculated as follows:

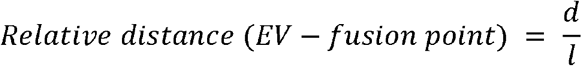

### Quantification of fusion defects

Formaldehyde-/methanol-fixed embryos were stained for luminal chitin. Chitin signal at DB (in stage 16 embryos) or DT (in stage 15 embryos) anastomoses was analyzed to determine whether the lumen was continuous or interrupted. The *y*^*1*^ *w*^*1118*^ strain was used as wild-type control.

### Quantification of Stac-MVBs

Confocal Z-stacks of tracheal metameres 4-6 were acquired in living embryos expressing EGFP-Stac in tracheal cells under the control of *btl-Gal4*. Stac-LROs were segmented in 3D by applying a smooth function, enhancing contrast (0.01 % saturated pixels) and using a manually defined threshold in Fiji. The number and volume of the vesicles was determined using the 3D object counter plugin in Fiji (Bolte & Cordelières, 2006).

### Statistics

Sample size (n) was not predetermined using statistical methods, but was assessed by taking into account the variability of a given phenotype, determined by the standard deviation. Experiments were considered independent if the specimens analyzed were derived from different parental crosses. During experiments investigators were not blinded to allocation. Sample size (n) is indicated in the figure legends or graphs. Data was tested for normality using the Shapiro-Wilk test. When data was not normally distributed, the Wilcoxon rank-sum test (R standard package) was used. P values were corrected for multiple testing using the Bonferroni-Holm method (Holm, 1979). For normally distributed data, the t-test was used from the R standard package. Since the EV data consists of non-continuous counting data we applied a simulated chi-squared test as follows. For the number of EVs a contingency table was generated from which the following test-statistic was calculated:

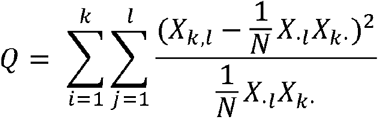

where N is the number of observations, *k* the number of different genotypes, and *l* the number of EV classes. *X*_*·l*_ and *X*_*k·*_ denote the column and row sum of the contingency table. The p-value was determined by generating n (10000) random permutations of the data and checking how many permutations yielded a larger Q statistic than the non-permutated data.

## Supplemental Figures

**Figure S1. Extracellular vesicles appear after dorsal trunk fusion and remain in close proximity to tracheal anastomosis sites**.

**(A**,**B)** Stills from time-lapse movies of living embryos (stage 13) expressing palm-mKate2 (magenta) and EGFP-Stac (green, A) or palm-mKate2 (magenta) and CD63-GFP (green, B) in tracheal cells after DT tube fusion (0 min). EGFP-Stac-positive EVs (A, arrowhead) and CD63-GFP-positive EVs (B, arrowhead) appeared after DT fusion and remained in close proximity to the DT anastomosis site. Tracheal lumen in (A, cross-section) is indicated by a dashed line. Stills in (B) are from Movie S2.

**(C)** Stills from time-lapse movie of embryo (stage 15) expressing EGFP-Stac in tracheal cells. Note that the Stac-positive EV (arrowhead) is largely immobile over 247 min and rapidly disappears during subsequent luminal liquid clearance (261 min). (C’) Kymograph of EGFP-Stac signal within the area indicated by a dashed rectangle in (C). Stills are from Movie S3. Scale bars: 10 µm.

**Figure S2. Stac localizes to intraluminal vesicles in secondary cells and in exosomes in the accessory gland lumen**.

**(A)** Scheme illustrating male reproductive male tract with accessory gland comprising main cells (light blue) and secondary cells (green). Secondary cells contain large endolysosomal vacuoles with ILVs, which are secreted as exosomes into the accessory gland lumen. Nuclei of the binucleate main and secondary cells are shown in cyan.

**(B-E)** Accessory glands of 3-day-old virgin male expressing EGFP-Stac (green) in secondary cells under the control of *Abd-B*-Gal4. Cell membranes are labeled with MM4-64 (magenta) and nuclei with Hoechst 33342 (cyan). (C) Accessory gland tip with secondary cells marked by EGFP-Stac expression (green). (D) Close-up of a single secondary cell shows that EGFP-Stac accumulates in ILVs (arrowhead) inside large MVBs. (E) Cross-section of the accessory gland tip (single plane, Z-depth 23.5 µm). Note that Stac-EVs (arrowhead) are detectable inside the accessory gland lumen, which also contains membranous microcarriers (Wainwright *et al*., 2021) labeled by MM4-64. (E’) shows a close-up of the region indicated in (E). Scale bars: (B) 500 µm; (C) 100 µm; (D, E, E’) 10 µm.

**Figure S3. Stac-MVBs localize between invading lumina and the FC-FC interface**.

**(A)** Scheme illustrating the localization of Stac-MVBs (green) in tracheal fusion cells (FCs). Adherens junctions (magenta) between fusion cells and the invading stalk cell (SC) lumina, and between FCs and the central lumen at the FC-FC interface are indicated. Tracheal cells are shown in petrol and the tracheal lumen in yellow.

**(B)** Dorsal branch fusion cells in embryo (stage 15) expressing endogenously tagged E-Cad::3xmTagRFP (magenta) and EGFP-Stac (green) in tracheal cells. (B’) shows a time series of the region marked in (B). Time is indicated. Arrowheads indicate Stac-positive MVBs that appear to merge with the central lumen at t=40 s. In single-channel images, EGFP-Stac intensities are displayed as a heat map (range indicated in (B)). Images were processed using Leica Lightning deconvolution in adaptive mode. The scheme to the right illustrates the position of EGFP-Stac MVBs (green) and E-Cad-labeled adherens junctions (magenta).

**(C)** Intensity profiles of EGFP-Stac and E-Cad::3xmTagRFP along the dashed line in (B’).

**(D)** Dorsal branch fusion cells in embryo (stage 15) expressing PLC*δ*-PH-EGFP (green) and mCherry-Stac (magenta) in tracheal cells. (D’) shows a single plane of the region marked in (D). Note that mCherry-Stac-positive MVBs accumulate between the invading lumina and the FC-FC interface.

Scale bars: (B, D) 5 µm; (B’) 1 µm; (D’) 2 µm.

**Figure S4. Identification of Rab GTPases involved in Stac-MVB formation**.

**(A)** Lateral view of dorsal trunk metameres 4-6 in living embryos (stage 15) expressing EGFP-Stac and Arl3 in tracheal cells under the control of *btl-Gal4*. Embryos carry a mutation in the indicated *Rab* locus. In case of X-chromosomal *Rab* loci (*Rab10, 18, 21, 27, 35, 39, 40*) embryos are either heterozygous or hemizygous for the mutation. For the remaining (autosomal) *Rab* loci, embryos are heterozygous for the mutation. *y w* was used as wild-type control (first panel). Note that EGFP-Stac-MVBs are ectopically induced throughout the tracheal system upon Arl3 misexpression. Embryos were scored for modification of the number of EGFP-Stac-MVBs by the *Rab* mutations.

**(B)** Quantification of Stac-MVBs in metameres 4 to 6 of embryos hetero- or hemizygous for the indicated *Rab* mutation. Stac-MVBs were quantified in 20 embryos per genotype. Wilcoxon rank-sum test: * p < 0.05.

**Figure S5. Model for extracellular vesicle release by tracheal cells**.

**(A)** Tracheal cells release Stac-positive (green) and Stac-negative EVs (grey). Tracheal lumen is shown in yellow, tracheal trunk cells in grey and tracheal fusion cells (FCs) in green. DT, dorsal trunk; DB, dorsal branch; TC, transverse connective. **(B)** Model for release of Stac-positive EVs by FCs (green) during tracheal lumen fusion. FCs migrate towards each other (top). After establishing cell-cell contact, a central lumen is formed at the FC-FC interface. Stac-positive MVBs, the formation of which depends on the GTPases Arl3, Rab27 and Rab35, localize between the invading lumina (yellow) of neighboring tracheal cells (grey) and the central lumen at the FC-FC interface. Stac protein (blue rectangles) localizes at the outer membrane of MVBs and inside intraluminal vesicles (ILVs; turquoise). Stac-containing EVs are released into the tracheal lumen upon fusion of Stac-MVBs with the luminal plasma membrane in FCs. ILVs may fuse with the outer membrane (black) of Stac-MVBs, providing an extra source of membrane material. **(C)** Stac-negative EVs are likely to be produced by cells throughout the tracheal tube and may include exosomes, derived from fusion of MVBs with the apical membrane of tracheal cells, as well as ectosomes, derived by shedding from the plasma membrane.

## Supplemental Movies

**Supplemental Movie 1**

EGFP-Stac-positive extracellular vesicles are present in the tracheal lumen. 3-D animation of the dorsal trunk lumen of an embryo expressing EGFP-Stac (green) and Verm-mRFP (magenta; labeling the lumen) in tracheal cells. Rendering of the tracheal lumen was performed in Imaris using the Surfaces function.

**Supplemental Movie 2**

Time-lapse movie of embryo expressing CD63-GFP (green) and palm-mKate2 (magenta) after DT fusion. A single confocal section containing a CD63-GFP-positive EV is shown. CD63-GFP-positive EVs appear after DT fusion and remain in close proximity to tube fusion sites.

**Supplemental Movie 3**

Time-lapse movie of embryo (stage 15) expressing EGFP-Stac (green) and palm-mKate2 (magenta) in tracheal cells. A single confocal section containing a EGFP-Stac-positive EV is shown. Note that EGFP-Stac EVs are immobile inside the tracheal lumen, but start to move and subsequently disappear rapidly during luminal liquid clearance (beginning at t=240 minutes).

