## Supplementary figures and images for "Tracheal tube fusion in *Drosophila* involves release of extracellular vesicles from multivesicular bodies"

### Figure S1

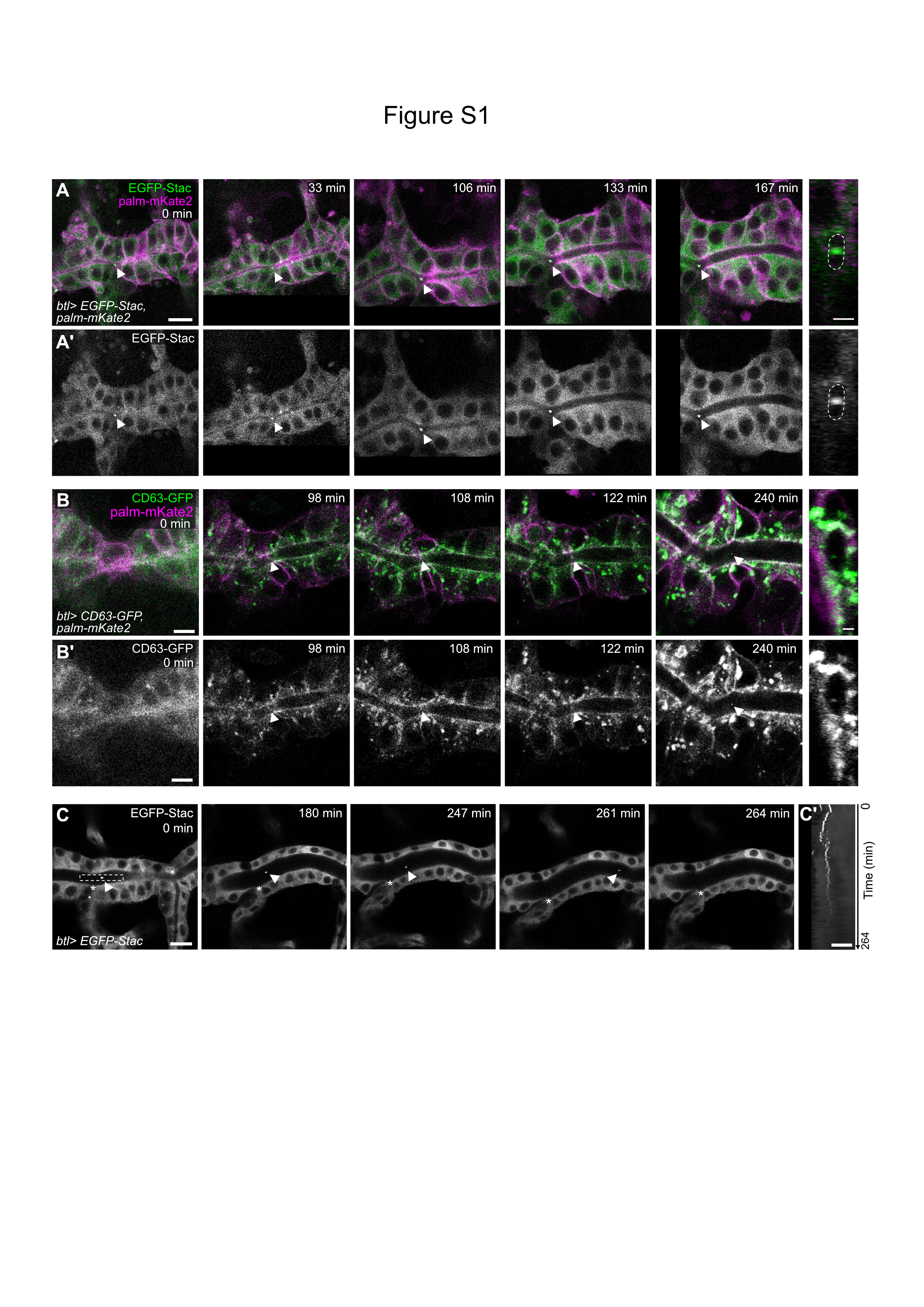

### Figure S2

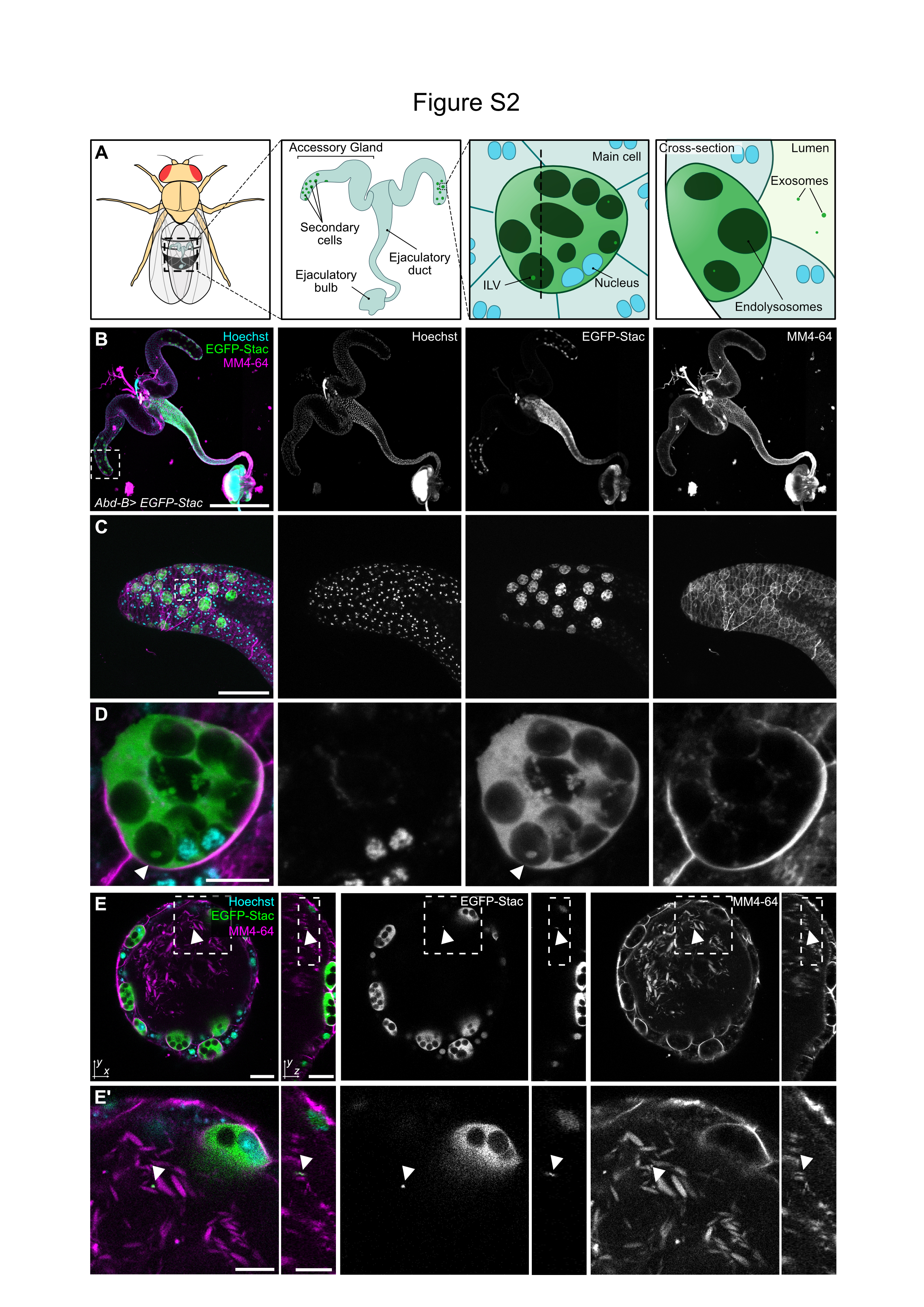

### Figure S3

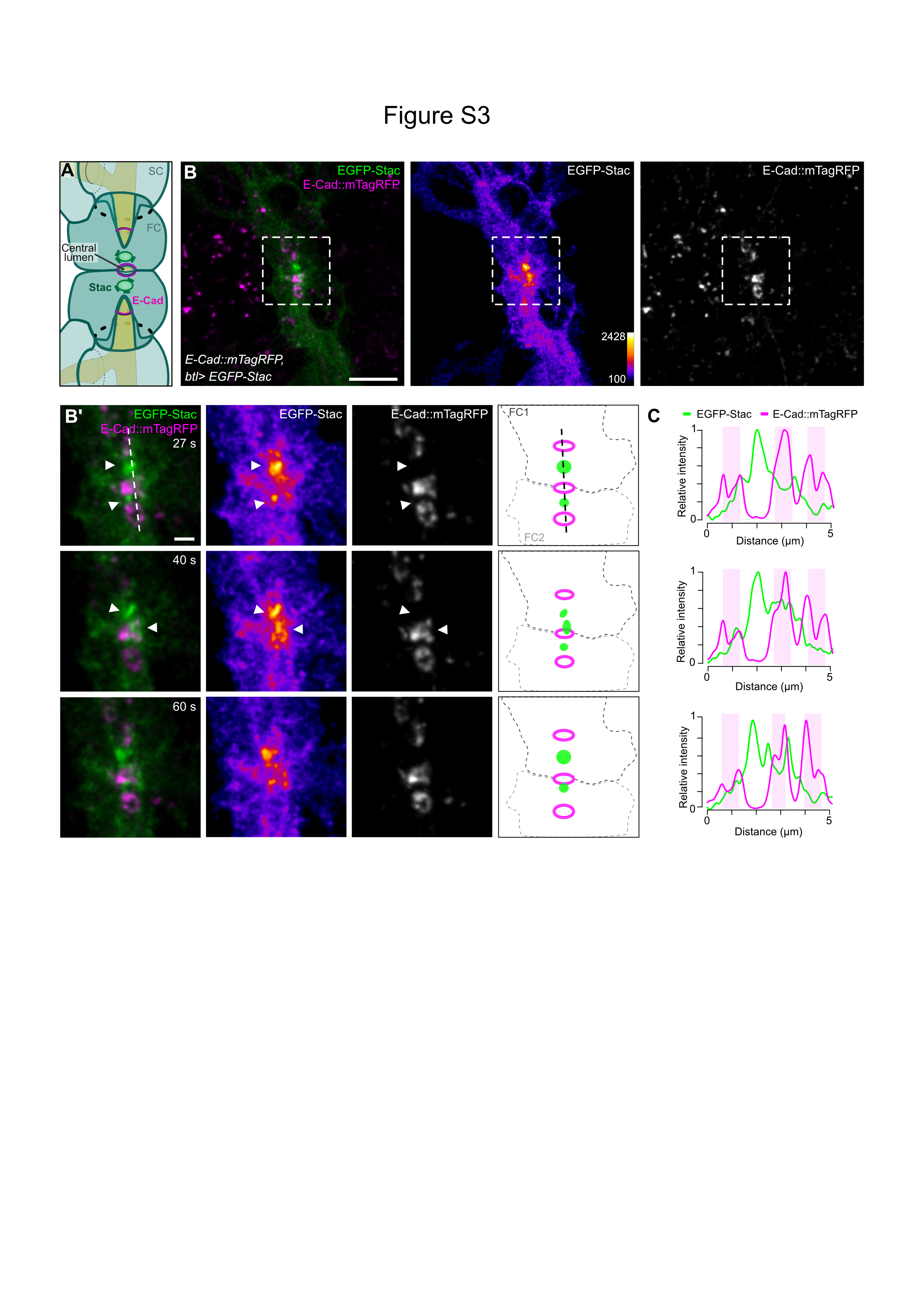

### Figure S4

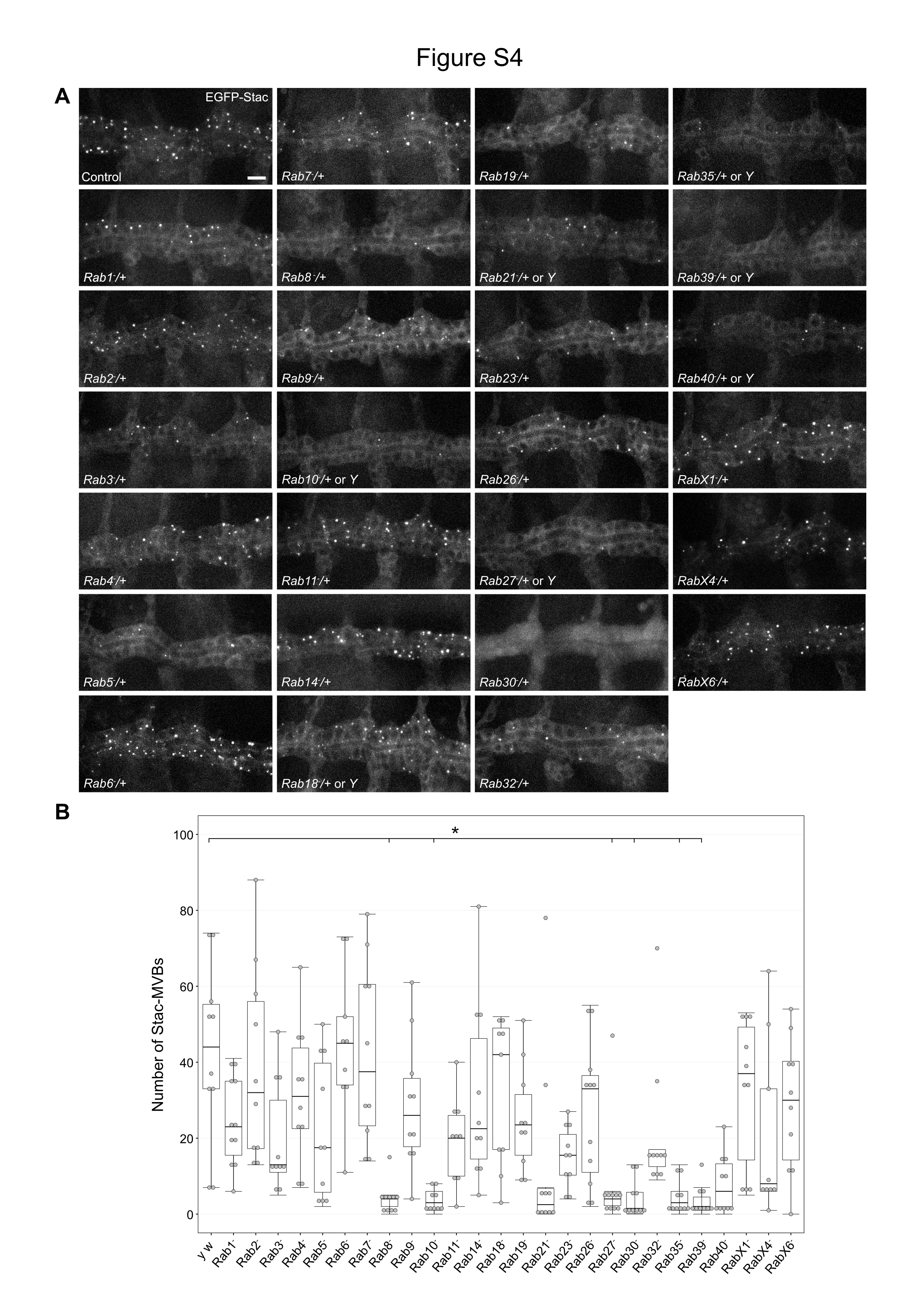

### Figure S5

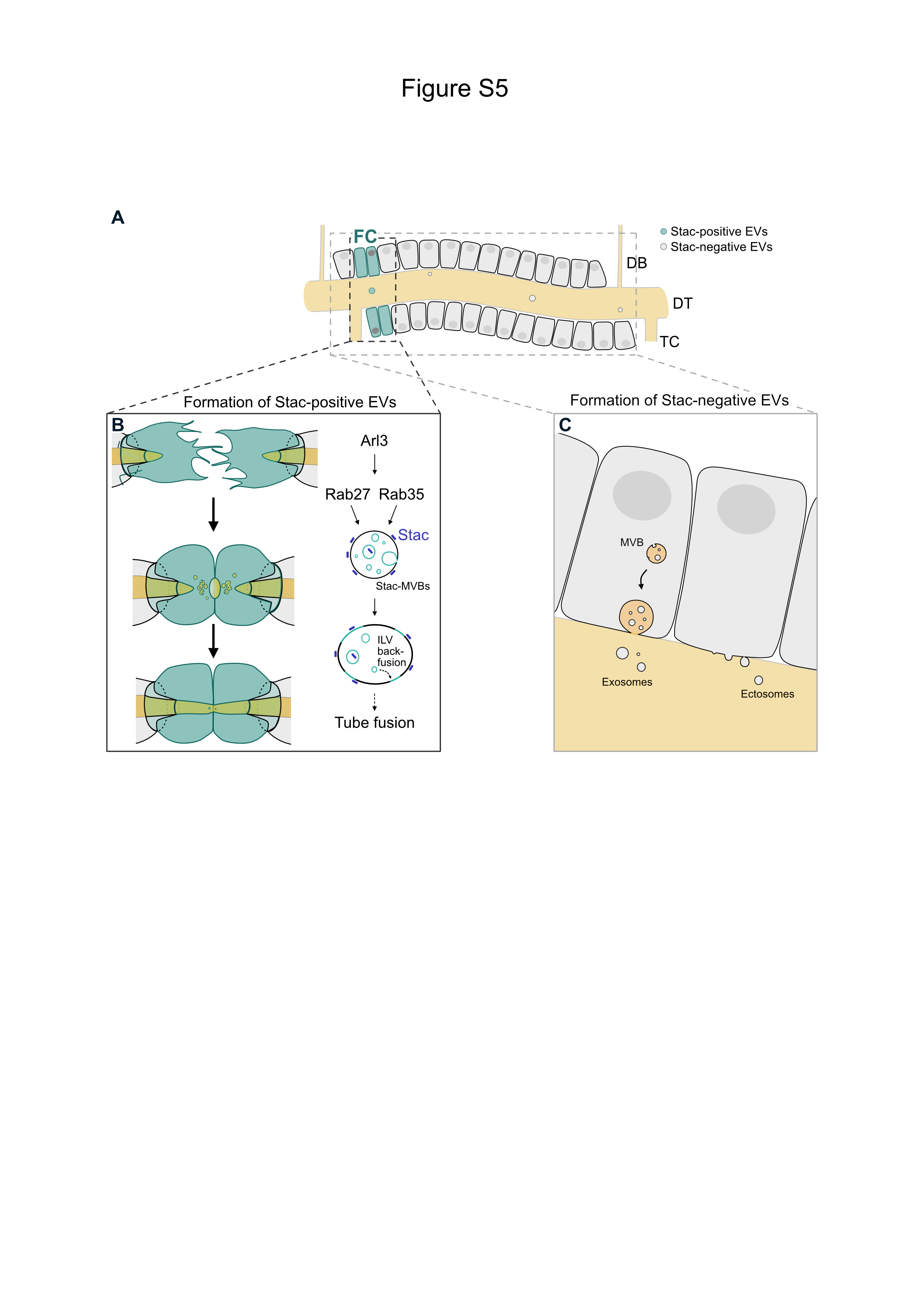
